# Activation of innate immunity selectively compromises mitochondrial complex I, proline oxidation and flight activity in the major arbovirus vector *Aedes aegypti*

**DOI:** 10.1101/2023.10.20.563264

**Authors:** Alessandro Gaviraghi, Ana Beatriz F. Barletta, Thiago Luiz Alves E Silva, Matheus P. Oliveira, Marcos H.F. Sorgine, Marcus F. Oliveira

**Affiliations:** Laboratório de Bioquímica de Resposta ao Estresse, Instituto de Bioquímica Médica Leopoldo de Meis, Universidade Federal do Rio de Janeiro, Rio de Janeiro, RJ 21941-590, Brazil; Instituto Nacional de Ciência e Tecnologia em Entomologia Molecular (INCT-EM), Rio de Janeiro, RJ 21941-590, Brazil; Laboratório de Bioquímica de Artrópodes Hematófagos, Instituto de Bioquímica Médica Leopoldo de Meis, Programa de Biologia Molecular e Biotecnologia, Universidade Federal do Rio de Janeiro, RJ 21941-590, Brazil; ABFB and TLAS: Laboratory of Malaria and Vector Research, National Institute of Allergy and Infectious Diseases, National Institutes of Health, Rockville, MD 20852, USA; MPO: Capacity Bio, Inc., Magnify at CNSI, CA 90095, USA

**Keywords:** Metabolism, proline, Zika, Dengue, bioenergetics, respiration, muscle, immunity, fitness, dispersal, flight, transmission

## Abstract

*Aedes aegypti* females are natural vectors of important arboviruses such as Dengue, Zika, and yellow fever. Mosquitoes activate innate immune response signaling pathways upon infection, as a resistance mechanism to fight pathogens and limit their propagation. Despite the beneficial effects of immune activation for insect vectors, phenotypic costs ultimately affect their fitness. However, the underlying mechanisms that mediate these fitness costs remain poorly understood. Given the high energy required to mount a proper immune response, we hypothesized that systemic activation of innate immunity would impair flight muscle mitochondrial function, compromising tissue energy demand and flight activity. Here, we investigated the dynamic effects of activation of innate immunity by intra-thoracic zymosan injection on *A. aegypti* flight muscle mitochondrial metabolism. Zymosan injection significantly increased defensin expression in fat bodies in a time-dependent manner that compromised flight activity. Although oxidant levels in flight muscle were hardly altered, ATP-linked respiratory rates driven by mitochondrial pyruvate+proline oxidation were significantly reduced at 24h upon zymosan injection. Oxidative phosphorylation coupling was preserved regardless of innate immune response activation along 24h. Importantly, rotenone-sensitive respiration and complex I-III activity were specifically reduced 24h upon zymosan injection. Also, loss of complex I activity compromised ATP-linked and maximal respiratory rates mediated by mitochondrial proline oxidation. Finally, the magnitude of innate immune response activation negatively correlated with respiratory rates, regardless of the metabolic states. Collectively, we demonstrate that activation of innate immunity is strongly associated with reduced flight muscle complex I activity with direct consequences to mitochondrial proline oxidation and flight activity. Remarkably, our results indicate a trade-off between dispersal and immunity exists in an insect vector, underscoring the potential consequences of disrupted flight muscle mitochondrial energy metabolism to arbovirus transmission.

## 1. Introduction

Resistance against infectious challenges requires the activation of immune mechanisms to target and eliminate pathogens, aiming at to restore host homeostasis and avoid premature death. To accomplish this, mammalians engage the so-called innate and adaptive immunity mechanisms that work in concert to generate a pro-inflammatory state that is absolutely required to cope with infection (Iwasaki and Medzhitov, 2015; Medzhitov, 2008). The innate immune system represents one of the first defensive barriers against invaders and comprises an array of soluble and cellular factors that immediately recognize pathogen-associated molecular patterns (PAMP) and subsequently mediate pathogen killing (Akira et al., 2001).

The challenges to mount an effective immune response are not limited to mammals. Despite the lower complexity, specificity, and the absence of an immunological memory, innate immune signaling pathways are remarkably conserved through evolution, acting as effective barriers to pathogens in insects (Galenza and Foley, 2019). The multi-task effort to mount a proper immune response in insects involves distinct tissues including *i)* the fat body, that controls systemic energy homeostasis and antimicrobial peptide (AMP) production (Arrese and Soulages, 2010)*; ii)* the hemocytes that mediate melanization, encapsulation and phagocytosis of pathogens (Lavine and Strand, 2002); *iii)* the midgut that represents the physical barrier to separate the external and internal milieu and also mediates the cross-talk between microbiota, internal organs and pathogens (Barletta et al., 2017; Das De et al., 2018). Regardless of the organ, key signaling pathways mediate innate immune responses, including the IMD and Toll pathways, orthologs of mammalian Tumor Necrosis Factor-alpha and Toll-Like Receptor, respectively, which results in production of the AMP (Barletta et al., 2017, 2012; Blandin et al., 2008; Imler et al., 2000; Keene et al., 2004; Sánchez-Vargas et al., 2009). While the IMD pathway is present in several insect tissues and recognizes mostly peptidoglycan from Gram-negative bacteria (Kaneko and Silverman, 2005), the Toll pathway essentially operates in the fat body and hemocytes, responding to Gram-positive bacteria or fungi PAMP (El Chamy et al., 2008). Beyond these mechanisms, insulin-like peptides and Target Of Rapamycin (TOR) as well as Akt, FOXO and AMP, complement the array of innate immune response mechanisms in insects (Buchon et al., 2014; Galenza and Foley, 2019; Myllymäki et al., 2014; Valanne et al., 2011).

Although the acute elimination of infectious agents by innate immune activation has a clear beneficial output on survival, several other life-history traits are impacted when inflammation turns chronic and uncontrolled (Akira et al., 2001; Bonneaud et al., 2003; Coussens and Werb, 2002; Martin et al., 2003). These include physiological, molecular, and metabolic alterations that ultimately compromise homeostasis and survival rates (Calvano et al., 2005; Franceschi et al., 2018). Indeed, an efficient immune response is energetically costly and poses critical trade-offs among nutritionally competing physiological processes such as reproduction, cognition, and motor activity (Rivera et al., 1998; Tidball, 2005). Hereof, evidence converges to the fact that a global shift in nutrient utilization from anabolism to catabolism is required in inflammatory states. In many cases, energy metabolism pathways are modified in multiple ways by the innate immune response, including: *i)* transient insulin resistance (Rera et al., 2012; Shi et al., 2006); *ii)* reduction in energy storage (Dionne et al., 2006; Rera et al., 2012)*; iii)* increased protein catabolism (Biolo et al., 1997); and *iv)* altered mitochondrial function (Brealey et al., 2002; Japiassú et al., 2011; Singer, 2014). These metabolic changes seem in principle beneficial, as they allow nutrient redistribution from high energy-demanding tissues to immune cell function. However, sustained catabolism and altered mitochondrial function have potentially harmful consequences for organisms (Brealey et al., 2002; Fouque et al., 2008).

Mitochondria are central organelles which play a multitude of critical cellular functions including energy homeostasis, redox balance, cell signaling, stress responses and beyond (Brand and Nicholls, 2011; Figueira et al., 2013; Friedman and Nunnari, 2014; Giacomello et al., 2020; Merry and Ristow, 2016; Vercesi et al., 2018). Cellular energy balance in aerobic organisms strongly depends on the ability of mitochondria to oxidize multiple substrates including carbohydrates, lipids and amino acids to allow ATP production by means of oxidative phosphorylation (OXPHOS). The electron flow through the mitochondrial redox complexes flows to ubiquinone through distinct dehydrogenases, including NADH dehydrogenase (complex I), proline dehydrogenase and glycerol phosphate dehydrogenase. Electrons are then transferred to complex III, cytochrome *c*, cytochrome *c* oxidase and finally to O_2_. The energy released by electron transfer is partially conserved as a protonmotive force, which is consumed by the F_1_F_o_ ATP synthase complex to allow ATP production. A general description of the reactions carried out by mitochondrial electron transfer system (ETS) complexes to transduce energy for ATP synthesis in *A. aegypti* is outlined in Figure 1.

**Figure 1.**
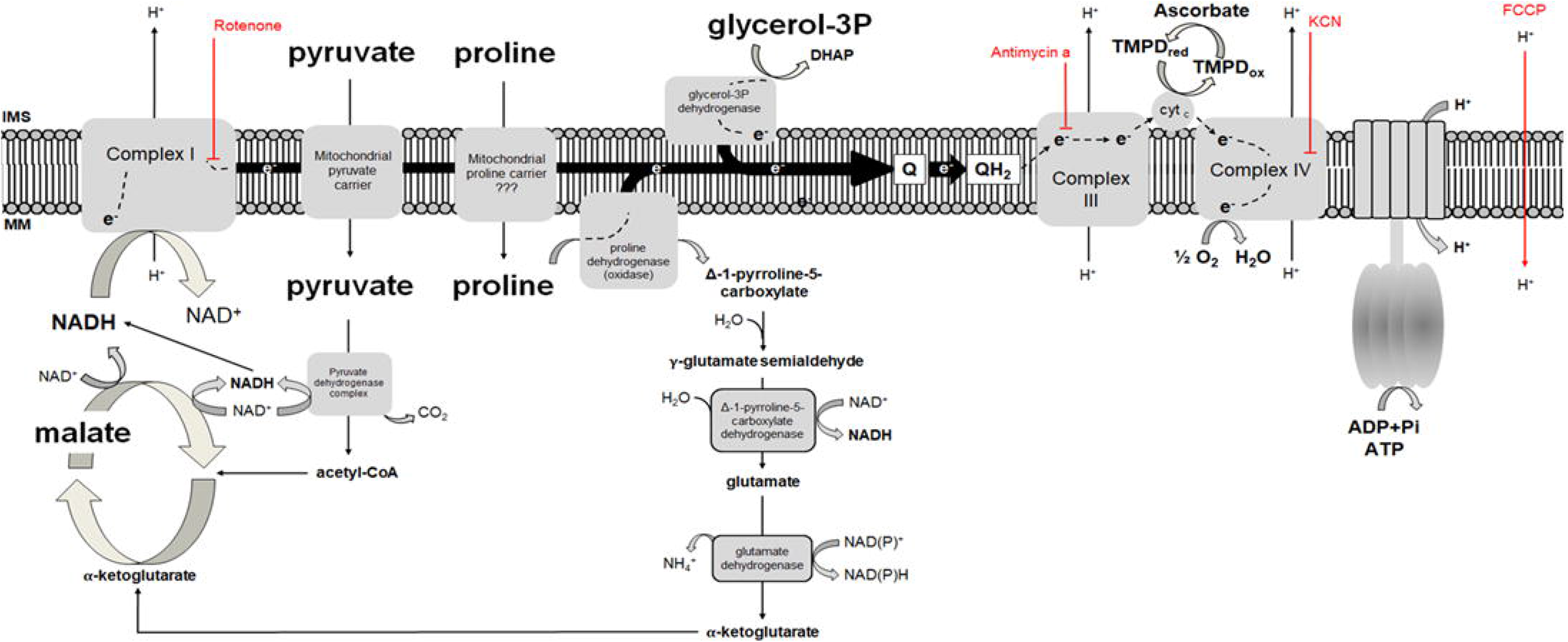
Schematic representation of the mechanisms involved in oxidative phosphorylation (OXPHOS). The action sites of pharmacological modulators are depicted in red and the different substrates oxidized by this organelle are shown in bold fonts. IMS: intermembrane space; MM: mitochondrial matrix; Q: oxidized ubiquinone; QH_2_: reduced ubiquinone. TMPD and ascorbate are substrates to specifically assess cytochrome *c* oxidase activity. Modified from (Soares et al., 2015).

Given the multitude of functions, it is not surprising that mitochondria directly participate in immunity, playing a key role in antiviral and antibacterial signaling, and in sterile cellular damage responses (Rambold and Pearce, 2018; West et al., 2011; Zhang et al., 2010). Conversely, during inflammatory challenges, mitochondrial structure, function, and life cycle are targeted, which ultimately impact cellular metabolism and viability (Buck et al., 2016; Calvano et al., 2005). Considering an extreme inflammatory insult, such as in sepsis, mitochondrial structure, function, and content are dramatically changed as a consequence of reduced tissue perfusion, excessive oxidants production, cytokines, as well as altered hormonal and gene expression (Cloonan and Choi, 2013; Rambold and Pearce, 2018; Stanzani et al., 2019). Indeed, the energy costs posed by the activation of the innate immune response are also manifested in insects, which are naturally exposed to a wide array of habitats and pathogens in nature. Interestingly, several metabolic consequences in insects were observed as a result of innate immune activation including *i)* increased nutrient intake to compensate for reduced energy availability (Moret and Schmid-Hempel, 2000)*; ii)* mitochondrial depolarization, loss of cell viability and reduced longevity by overexpression of AMP (Badinloo et al., 2018); and *iii)* mitochondrial damage and extrusion from enterocytes by bacterial hemolysin (Lee et al., 2016). Metabolic impacts caused by mosquito infection were also reported including dynamic alterations in mitochondrial O_2_ consumption without affecting cellular ATP levels (Santana-Román et al., 2021). Also, refractoriness to *Plasmodium* infection is associated with strong reductions in OXPHOS in *Anopheles gambiae* mosquitoes (Oliveira et al., 2011). Large-scale quantitative proteomic studies during arboviral infection have shown remarkable changes in the expression of key metabolic enzymes including those involved in glycolysis, tricarboxylic acid cycle, and OXPHOS (Martins et al., 2021; Vasconcellos et al., 2022, 2020). However, direct evidence demonstrating alterations on insect mitochondrial function resulting from immune activation remains largely obscure.

Since the activation of innate immune response represents a competing process for the physiological energy demands, we postulated that a trade-off would exist between systemic immunity and flight activity in an insect vector, as a consequence of disrupted mitochondrial energy metabolism in flight muscle. Here, we investigated the effects of activation of innate immunity by intra-thoracic injection of zymosan on *A. aegypti* flight muscle mitochondrial metabolism. We demonstrate that activation of innate immunity has detrimental effects on mitochondrial physiology due to selective reduction in complex I activity, which ultimately compromise OXPHOS and energy supply to sustain flight activity. Remarkably, our results indicate that a trade-off between dispersal and immunity exists in *A. aegypti*, underscoring the potential consequences of disrupted flight muscle mitochondrial energy metabolism to arbovirus transmission.

## 2. Results

### 2.1. Zymosan activates *A. aegypti* innate immune response and impacts flight activity

Our first attempt was to establish a robust model of innate immune response activation in young adult *A. aegypti* females fed exclusively with sucrose. The experimental approach chosen was based on the injection of either an isotonic phosphate buffered solution (hereafter named PBS) or a 5 mg/mL zymosan A stock suspension (hereafter named Zymo) (Figure 2). Zymosan A is a cell wall component of fungi composed of β-glucan and mannan (Di Carlo and Fiore, 1958) and is used as a classical immune response activator in several models including insects (Barletta et al., 2012; Wang et al., 2015). Insects were injected through the thoracic cavity with a Hamilton syringe with 1 μL of PBS or Zymo and then several parameters were analyzed along 24 h post-injection. Figure 3A shows that both PBS and Zymo injections caused no apparent effects on survivorship for up to 7 days and also on insect body weight (Figure 3B).

**Figure 2.**
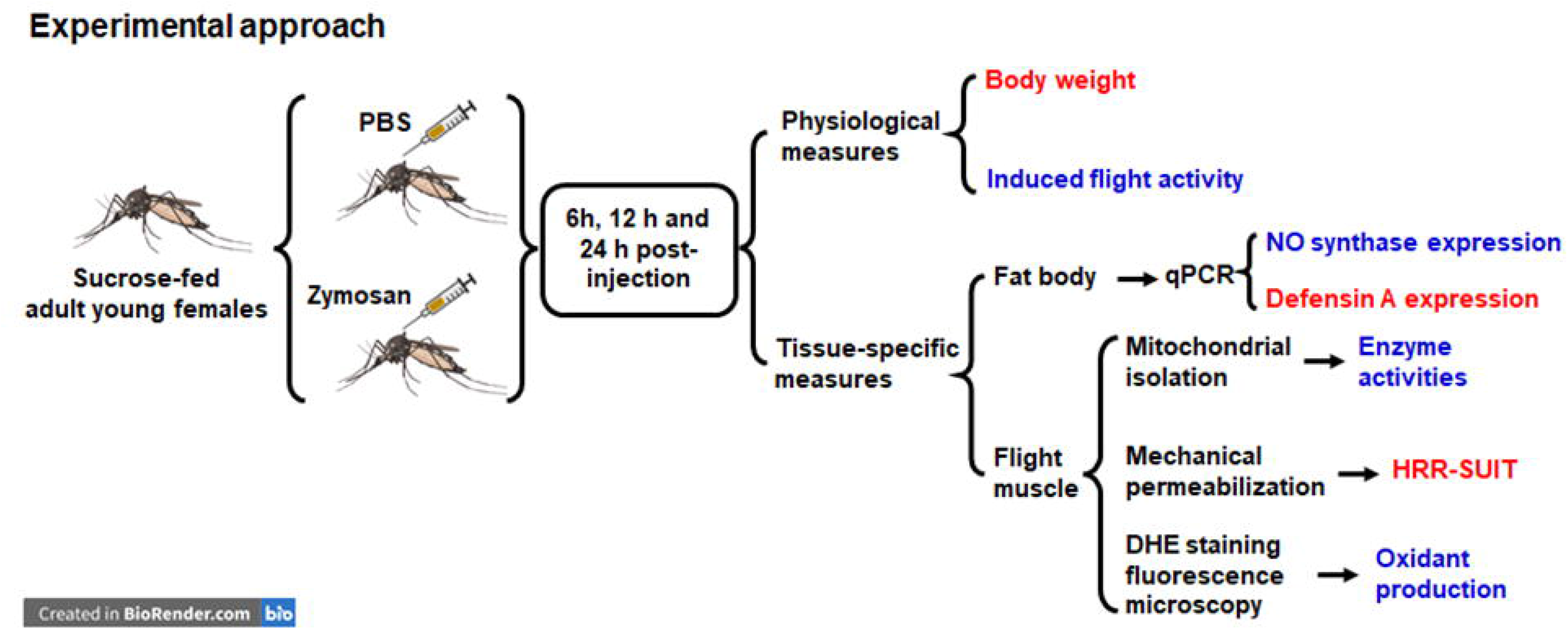
Experimental approach used in this study to assess the effects of innate immune response activation on *A. aegypti*. Adult young females fed exclusively on sucrose were injected with PBS or Zymosan and several physiological and tissue specific measures were determined at different timepoints. Red outputs represent those that were assessed from 6-24 h upon injection while blue ones only through 12-24 h.

**Figure 3.**
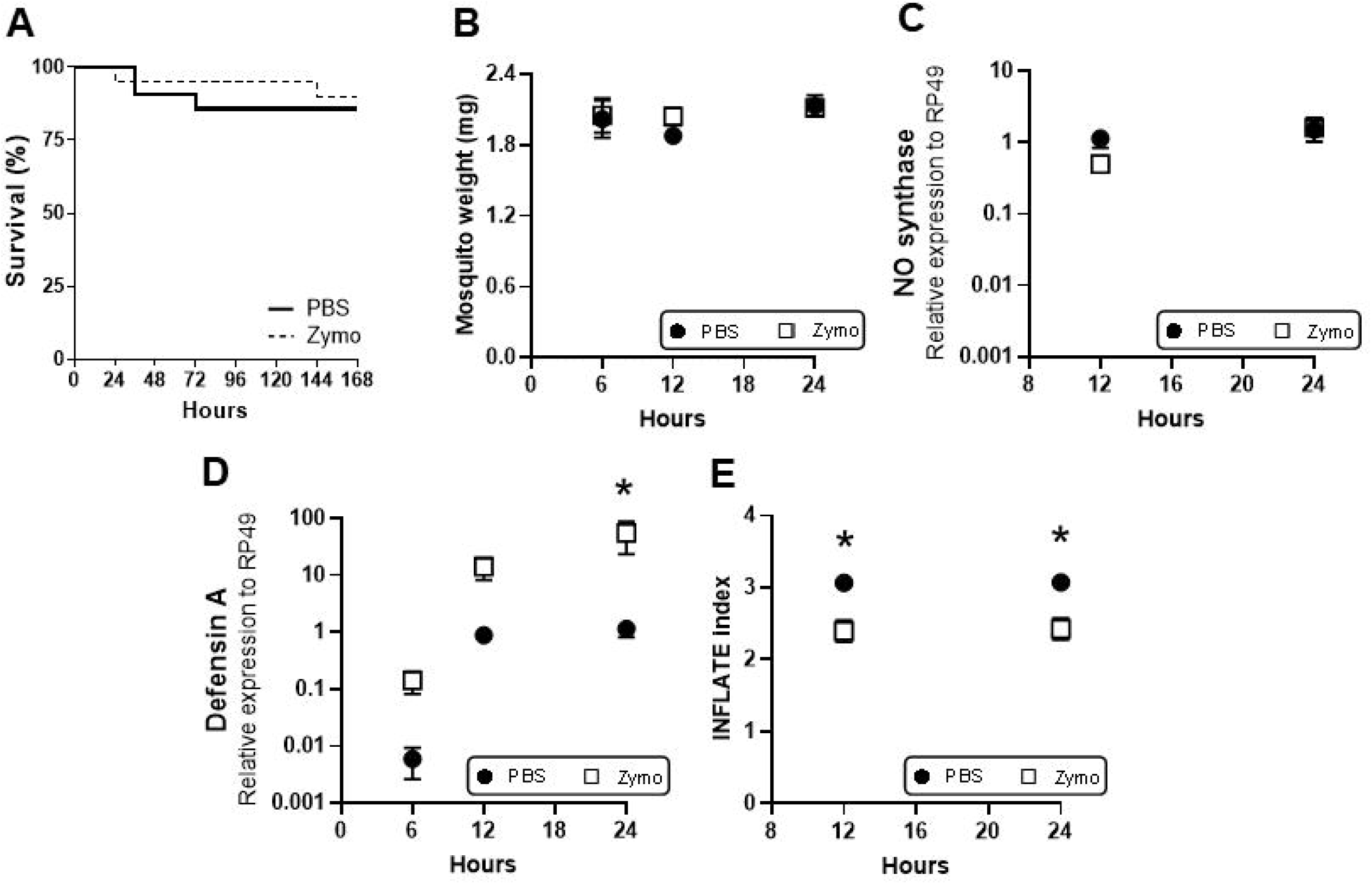
Zymosan potently activates *A. aegypti* immune response and compromise flight activity. (A) Representative survival curve of three different experiments, obtained after injection with PBS (solid line) or Zymo (dashed line). Each experiment was carried out using 30 mosquitoes for any group. (B) Mosquito weight was not affected after injection of either PBS (black circles), or Zymo (white squares), the insects were chilly-anesthetized and their total body weights were determined using an analytical balance with a precision of 0.1 µg. Nitric oxide synthase (C) and Defensin a (D) expression were determined in fat bodies of insects injected with either PBS (black circles, n=8-9) or Zymo (white squares, n=12) at 6, 12 and 24h. Comparisons between groups over time was done by using the Friedmańs and Durbin-Conover test (*p=0.04). (E) Flight activity was measured using the INFLATE method, as previously reported (Gaviraghi and Oliveira, 2020) in a group of 10 mosquitoes 12h and 24 h after injection with PBS (black circles, n=8) or Zymo (white squares, n=8). Comparisons between groups were done by 2-way ANOVA and *a posteriori* Šídák’s tests (F values of time, F_time_(1, 30) = 0.16, *p* = 0.692; F values of injection, F_inject_(1, 30) = 205.9, * *p* < 0.0001). Data are expressed as mean ± standard error of the mean (SEM) of at least four different experiments.

The activation of innate immune response was assessed through RT-qPCR by measuring the expression of the immune mediators nitric oxide synthase (NOS) and defensin A in fat bodies at 12 h and 24 h post-injection. We observed that expression of NOS was not affected by Zymo injection (Figure 3C), which strikingly contrasts with a potent upregulation of defensin A (Figure 3D). Indeed, the dynamics of immune activation reveal a time-dependent increase on defensin expression, starting as early as 6 h and reaching maximal levels at 24 h post-injection regardless of the treatment (PBS or Zymo). In this regard, we observed that injection of a sterile PBS solution caused an increase in defensin A expression, which was interpreted as a result of the simple injury caused by the injection itself. However, the exposure to Zymo elicited a powerful activation of the innate immune response as revealed by a remarkable increase in defensin A expression relative to PBS as early as 6 h post-injection (Figure 3D). However, significant differences were only observed at 24 h post-injection, where defensin A expression in Zymo-injected insects was about 45 times higher relative to PBS-injected ones. To check the potential side-effects of innate immune activation on *A. aegypti* physiology, we assessed the flight ability by using the induced flight activity test (INFLATE) developed in our laboratory (Gaviraghi and Oliveira, 2020). Figure 3E shows that Zymo injection significantly reduced the induced flight activity by 24 % as early as 12 h relative to PBS-injected insects. Therefore, Zymo injection in the thoracic cavity of *A. aegypti* activates innate immune response and reduces flight activity, without any apparent effects on body mass and survivorship. Since the dispersal of *A. aegypti* requires flight muscle activity, we then focused the next experiments on that tissue.

### 2.2. Preserved redox balance upon the activation of innate immune response

Reactive O_2_ (ROS) and nitrogen species (RNS) are known oxidants produced by physiological cell metabolism. However, the production of a variety of oxidant species, including hydrogen peroxide, superoxide, and nitric oxide, dramatically increases upon a traumatic or an infectious stimulus (Babior et al., 1973). The immune-mediated production of cellular oxidants is highly variable, depending not only on the magnitude, but also on the time and tissue where the insult takes place. Therefore, to investigate whether reduced flight ability in Zymo-injected mosquitoes was associated with redox imbalance in flight muscles, we assessed the levels of total oxidants by quantifying dihydroethidium (DHE) fluorescence in flight muscles. Figure 4 shows that potent activation of innate immune response observed 24 h upon Zymo injection in *A. aegypti* caused no apparent changes in total oxidant levels in flight muscle. Therefore, the activation of the innate immune response does not affect flight muscle redox balance.

**Figure 4.**
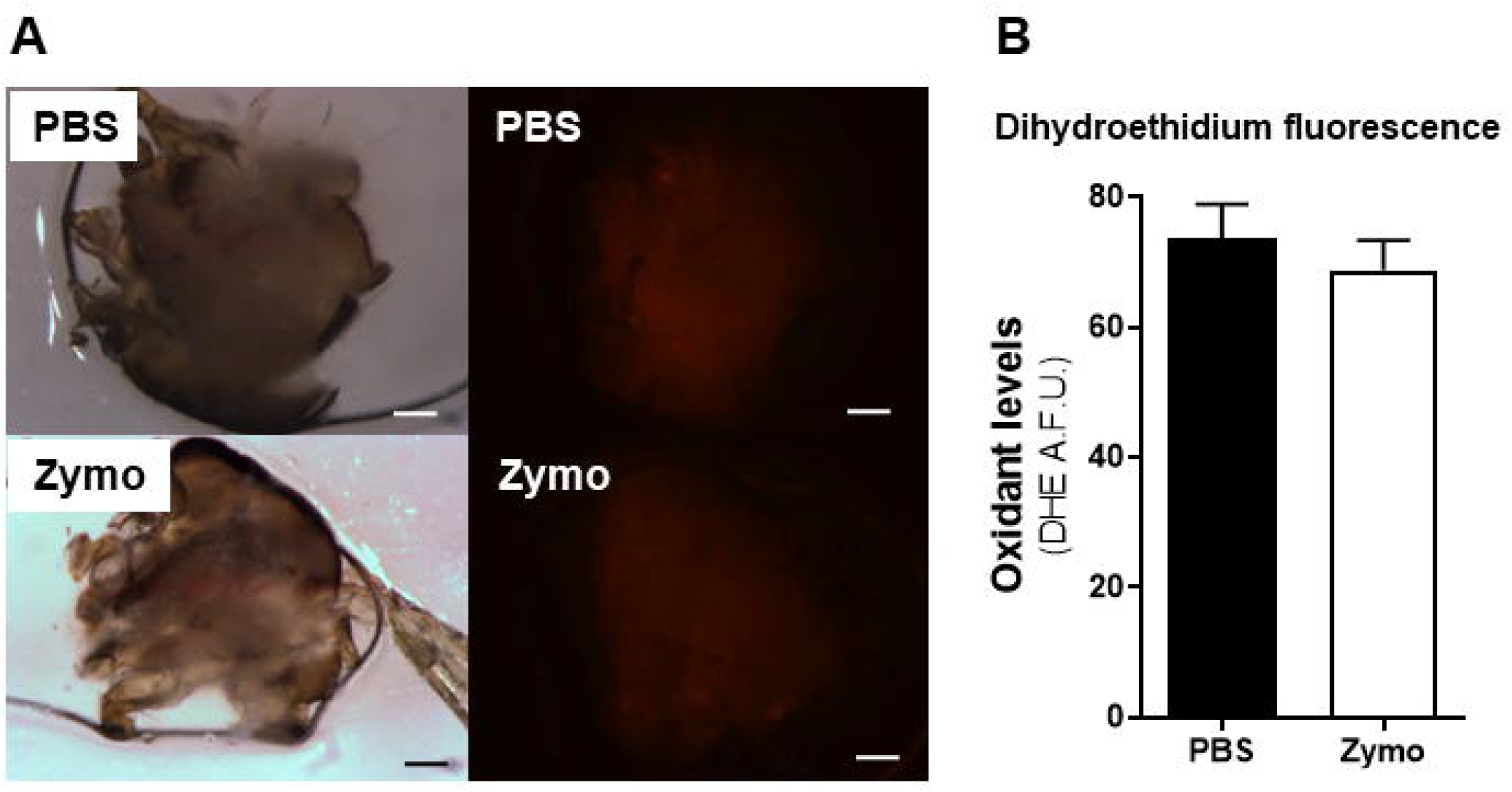
Activation of immune response does not change intracellular oxidants levels in flight muscle. (A) Representative images of hemi-thorax flight muscle of mosquitoes 24 h after injection with PBS or Zymo. Left are bright field images of thoraces and right are fluorescence stereomicroscopy of intracellular oxidants levels assessed by dihydroethidium (DHE) staining. Scale bars represent 200μm. (B) The average flight muscle DHE fluorescence intensity of PBS (black bar, n=6) and Zymo (white bar, n=8) injected insects. Data are expressed as mean ± standard error of the mean (SEM) of at least four different experiments for both PBS and Zymo groups.

### 2.3. Mitochondrial O_2_ consumption is strongly affected by innate immune response activation

We next examined the metabolic consequences of innate immune activation in flight muscles by assessing mitochondrial O_2_ consumption rates through high-resolution respirometry (HRR) coupled to substrate-uncoupler-inhibitor titration (SUIT) protocols using pyruvate and proline as fuels. The reason to use this combination lies on the fact that pyruvate and proline give the highest respiratory rates in *A. aegypti* flight muscles, as previously reported by our laboratory (Gaviraghi et al., 2019; Gaviraghi and Oliveira, 2019; Soares et al., 2015). To accomplish this, we utilized a method of mechanical permeabilization of a single *A. aegypti* thorax developed by our laboratory to study mitochondrial physiology *in situ* (Gaviraghi and Oliveira, 2019). This method allows the selective permeabilization of the plasma membrane and the assessment of flight muscle mitochondrial respiratory rates in a single mosquito, greatly enhancing our ability to study energy metabolism at individual level. Indeed, this procedure was extended and validated to study mitochondrial physiology in adult *Drosophila melanogaster* fruit flies (Gaviraghi et al., 2021). Figure 5B shows representative O_2_ consumption traces of mechanically permeabilized flight muscles from PBS (solid line) and Zymo (dashed line) injected insects 24 h upon immune activation. We observed that respiratory rates from Zymo-injected insects were in general lower than those from PBS-injected ones, suggesting reduced metabolic capacity of flight muscle mitochondria. We also observed that three out of the five mitochondrial metabolic states calculated, as depicted in Figure 5A, were significantly reduced upon Zymo injection. Despite leak respiratory rates were not significantly altered between groups over time, we observed a slight increase at 12 h in Zymo-injected insects suggesting a mild proton leak associated with immune activation (Figure 5C). Indeed, a significant effect of time (F_time_ *p* = 0.013) on leak respiratory rates was identified by 2-way ANOVA. Importantly, we observed that innate immune activation was associated with a time-dependent reduction in respiratory rates, especially at OXPHOS, spare and ETS respiratory rates (Figures 5D, 5E and 5F). Maximal inhibitory effects in these mitochondrial metabolic states were observed only at 24 h upon immune activation where respiratory rates of Zymo-injected insects were significantly lower than PBS-injected insects (Figures 5D, 5E, and 5F). We also observed proportional decreases in OXPHOS and ETS respiratory rates (17 and 23%, respectively) relative to their respective 6 h time points in Zymo-injected insects. Interestingly, significant reductions in spare respiratory rates were seen over time (F_time_ *p* = 0.035) regardless of the treatment suggesting that this metabolic state is much more sensitive to immune activation than any other mitochondrial functional parameter assessed (Figure 5E). However, the extent by which the ETS respiratory rates were affected upon immune activation was different among groups as in Zymo-injected insects a significant reduction in this parameter was observed at 24 h (Figure 5F). Also, in PBS-injected insects, the relative decline in ETS respiratory capacity was ∼30% at 12 h remaining stable afterward, while this metabolic state consistently reduced over time in Zymo-injected insects reaching inhibitions close to 52% at 24 h. Given that ETS respiratory rates reflect how close mitochondria operates to their bioenergetic limit, we postulate that reductions in this parameter upon the activation of innate immune response might be a consequence of a disrupted mitochondrial pyruvate/proline transport and/or oxidation rather than on ADP phosphorylation or mitochondrial inner membrane permeability.

**Figure 5.**
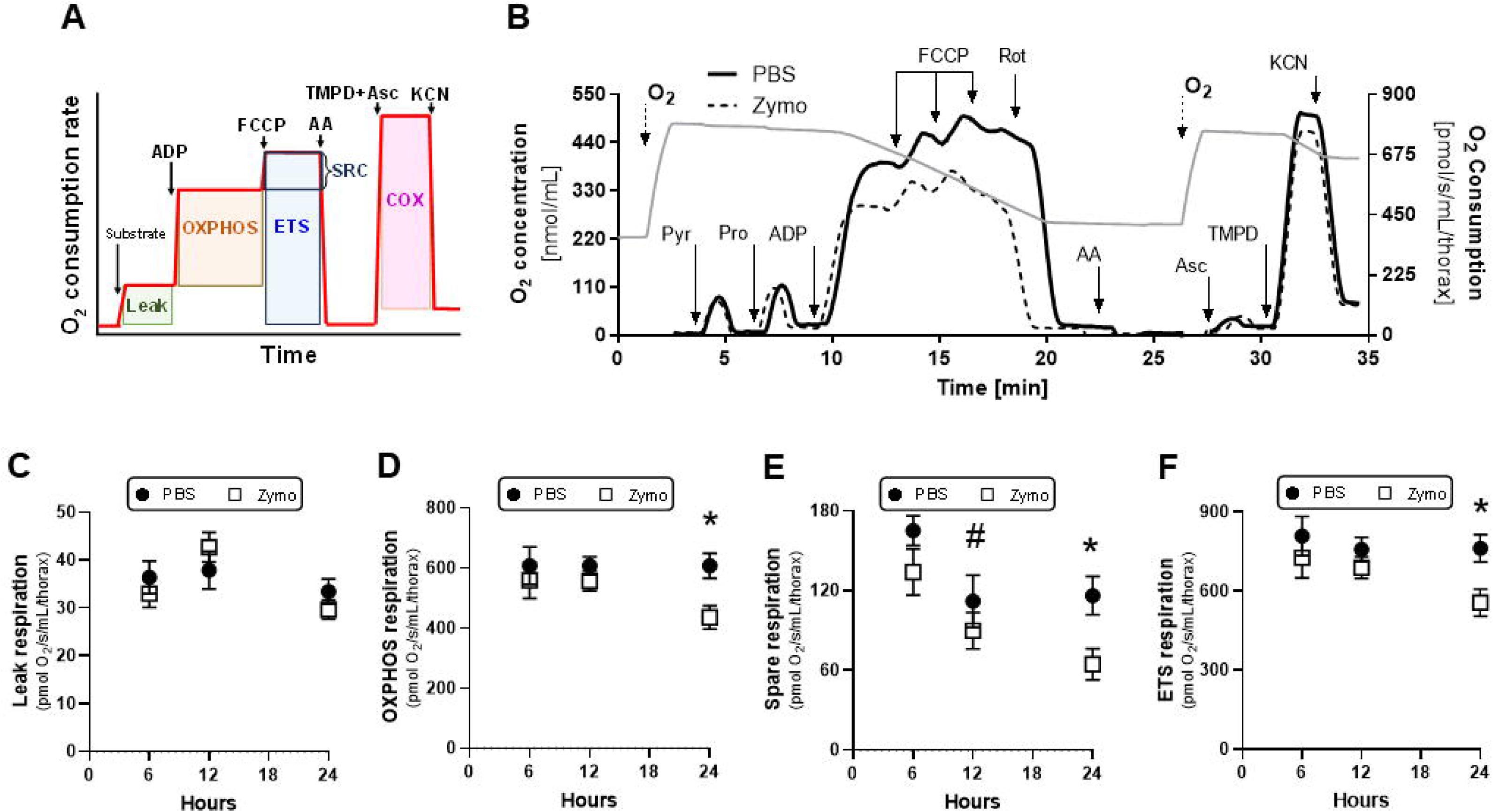
Activation of innate immune response compromise mitochondrial O_2_ consumption. (A) Schematic representation of a high resolution respirometry experiment. Red line represents a typical trace of O_2_ consumption rates in permeabilized cell or isolated mitochondria upon the addition of specific substrates and OXPHOS modulators. From these measures, five mitochondrial metabolic states were determined: i) the proton leak respiration (leak, green box) was assessed by the addition of oxidizable substrate and subtracting non-mitochondrial respiration generated by complex III inhibitor AA (grey box); ii) the ATP-linked respiration (OXPHOS, orange box) was derived by subtracting the proton leak respiration from ADP-induced O_2_ consumption; iii) the maximal respiratory rates (ETS, blue box) was determined upon the collapse of mitochondrial membrane potential by the addition of FCCP and subtracting non-mitochondrial respiration induced by the complex III inhibitor AA; iv) the spare respiratory capacity (SRC) was calculated by subtracting OXPHOS respiration from maximal respiratory capacity and v) the cytochrome c oxidase (COX, pink box) activity was determined by using TMPD+ascorbate as substrates and subtracting non-COX O_2_ consumption upon adding the COX inhibitor KCN. (B) Representative traces of O_2_ consumption rates (solid and dashed lines) and concentration (grey solid line) from mechanically permeabilized *A. aegypti* flight muscle using pyruvate+proline as substrates 24 h upon injection with PBS (solid line) or Zymo (dashed line). Arrows indicate the substrates and OXPHOS modulators added in the experiment as described in the methods section. (C-F) Quantification of flight muscle mitochondrial metabolic states of PBS (black circles) and Zymo (white squares) injected insects using 10 mM pyruvate-proline as substrates. (C) Leak respiration. Comparisons between groups were done by 2-way ANOVA and *a posteriori* Šídák’s tests. F_time_ (1.9, 43) = 5.0, *p* = 0.013; F_inject_ (1, 46) = 0.101, *p* = 0.752. (D) OXPHOS respiration. Comparisons between groups were done by 2-way ANOVA and *a posteriori* Šídák’s tests. F_time_ (1.96, 44.1) = 1.47, *p* = 0.24; F_inject_ (1, 45) = 5.99, *p* = 0.018. * A significant difference between PBS and Zymo was detected at 24 h (* *p* = 0.019). (E) Spare respiration. Comparisons between groups were done by 2-way ANOVA and *a posteriori* Šídák’s tests. F_time_ (1.96, 45.1) = 6.66, *p* = 0.0031; F_inject_ (1, 46) = 7.1, *p* = 0.011. * A significant difference between PBS and Zymo was detected at 24 h (* *p* = 0.035). (F) ETS respiration. Comparisons between groups were done by 2-way ANOVA and *a posteriori* Šídák’s tests. F_time_ (1.8, 41.2) = 1.8, *p* = 0.18; F_inject_ (1, 46) = 6.15, *p* = 0.017. * A significant difference between PBS and Zymo was detected at 24 h (* *p* = 0.031). Data are expressed as mean O_2_ consumption rates (pmol/s/mL/thorax) ± standard error of the mean (SEM) of at least seven different experiments.

### 2.4. Activation of innate immune response specifically affected complex I activity

In order to determine the mechanism by which activation of immune response reduces respiratory rates provided by pyruvate/proline metabolism, we assessed the activity of key mitochondrial enzymes by spectrophotometric quantification in isolated mitochondria, or polarographically by HRR-SUIT in mechanically permeabilized flight muscles. To determine the contribution of complex I to respiratory rates driven by pyruvate and proline oxidation, we determined the inhibitory effect of rotenone on uncoupled O_2_ consumption of permeabilized flight muscles from PBS and Zymo-injected insects at 6, 12 and 24 h post-injection. We observed that complex I-linked respiration was significantly reduced in Zymo-injected insects at 24 h, while no apparent changes were seen in earlier time points (Figure 6A). In order to certify this result, we spectrophotometrically assessed complex I activity in flight muscle isolated mitochondria by using an independent method of cytochrome *c* reduction assay. Figure 6B confirms that 24 h upon immune activation by Zymo caused a significant reduction in complex I activity. Considering the diversity of mitochondrial phenotypes generated by the activation of immune response (Brealey et al., 2004, 2002; Callahan and Supinski, 2005; Japiassú et al., 2011; Protti et al., 2007; Samavati et al., 2008; Vanasco et al., 2012), we then determined the activity of three critical mitochondrial enzyme activities: the glycerol 3 phosphate dehydrogenase (G3PDH), citrate synthase (CS) and the cytochrome *c* oxidase (COX). We observed that none of the three enzyme activities assessed were affected by Zymo injection strongly indicating that specific complex I dysfunction is a hallmark of immune activation in our experimental settings (Figures 6C-E). Given that G3PDH, CS and COX were not affected by Zymo, we suggest that reduced respiratory rates observed were not a consequence of lower mitochondrial content (Figure 6), but rather of compromising complex I activity.

**Figure 6.**
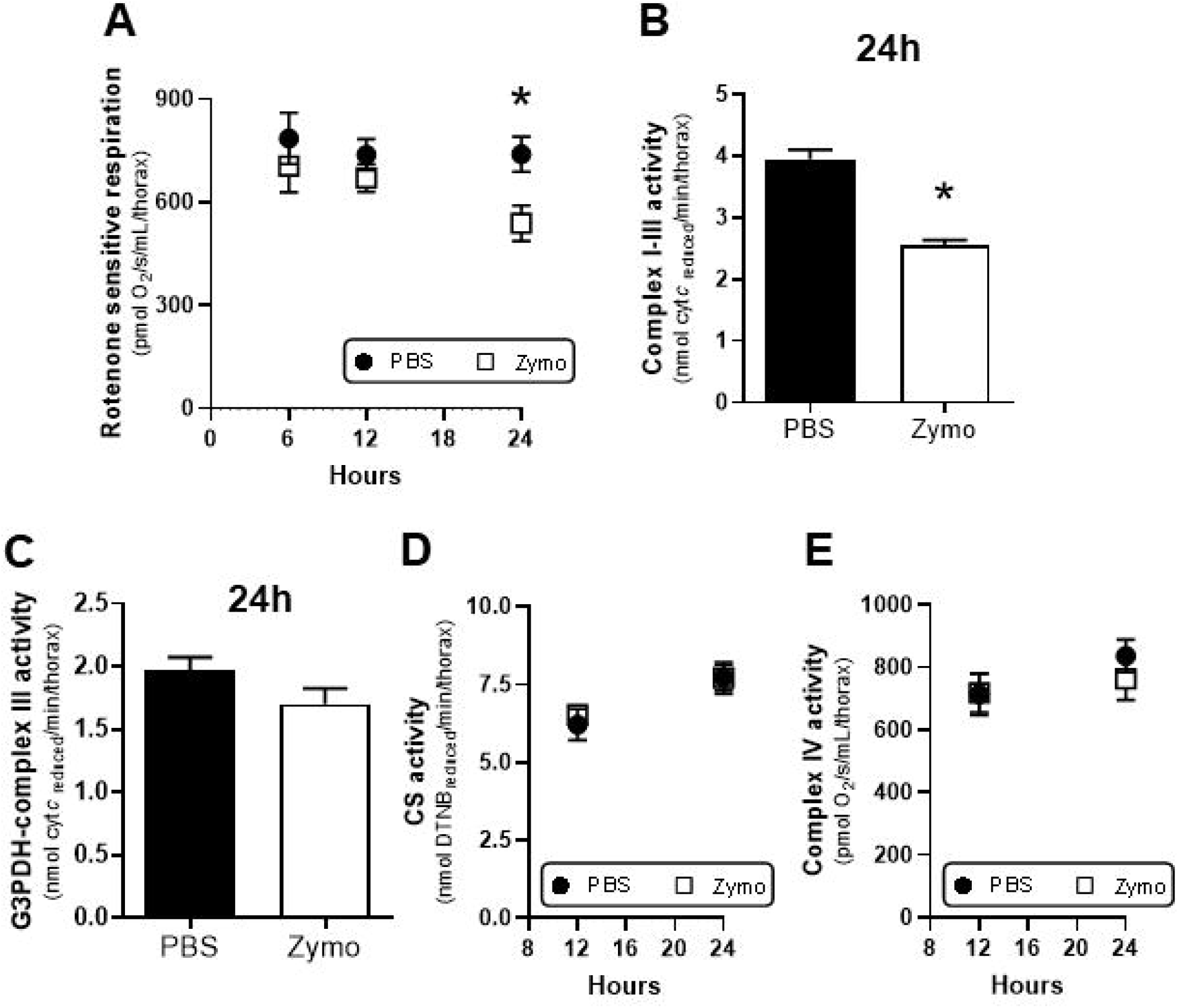
Specific reduction of complex I is a hallmark of immune activation. The activity of mitochondrial enzymes was determined spectrophotometrically in isolated mitochondria, or polarographically in mechanically permeabilized flight muscles. (A) Respiratory rates linked to complex I activity were quantified in flight muscles of PBS (black circles) and Zymo (white squares) injected insects by using 10 mM pyruvate-proline, 2 mM ADP and 2.5 µM FCCP following by the addition of 0.5 μM rotenone. Complex I-linked O_2_ consumption rates were calculated by subtracting the rotenone-insensitive O_2_ consumption rates by the FCCP-uncoupled respiration. Comparisons between groups were done by 2-way ANOVA and Šídák’s multiple comparisons test F_time_ (1.62, 37.30) = 1.79, *p* = 0.185 F_inject_ (1, 46) = 5.92, *p* = 0.02 relative to PBS 24h. (B) Complex I-III activity was assessed spectrophotometrically in flight muscles of PBS (black bar) and Zymo (white bar) exposed insects at 24h post injection by following the rates of cytochrome *c* reduction at 550 nm as described in the methods section. Comparison between groups were done by Mann Whitney with * *p* = 0.03 relative to PBS. (C) G3PDH-complex III activity was assessed spectrophotometrically in flight muscles of PBS (black bar) and Zymo (white bar) exposed insects at 24 h post injection by following the rates of cytochrome *c* reduction at 550 nm as described in the methods section. (D) Citrate synthase activity was assessed spectrophotometrically in flight muscles of PBS (black circles) and Zymo (white sqaures) exposed insects at 12 and 24 h post injection by following the rates of DTNB reduction at 412 nm as described in the methods section. (E) Cytochrome *c* oxidase (COX) activity was assessed by respirometry in flight muscles of PBS (black circles) and Zymo (white sqaures) exposed insects at 12 and 24 h post injection by following the O_2_ consumption rates as described in the methods section. Data are expressed as mean ± standard error of the mean (SEM) of at least four different experiments for both PBS and Zymo groups.

### 2.5. OXPHOS coupling efficiency is preserved despite reduced complex I activity

To determine whether loss of complex I activity by the innate immune response would compromise the energy coupling between respiration and ATP synthesis through the OXPHOS, we calculated the respiratory control ratios (RCRs) and flux control ratios (FCR) of permeabilized flight muscles from PBS and Zymo-injected insects from 6 to 24 h upon injection. The RCR values are empirical measures which reflect the integrity of the mitochondrial inner membrane and are defined as the ratio between ADP-stimulated/leak respiratory rates (Brand and Nicholls, 2011). The RCR values strongly vary from fully coupled (infinity values) to fully uncoupled mitochondria (1.0 values) and depend on the oxidized substrate and the magnitude of the inner membrane proton permeability. We observed in our experimental settings that innate immune activation caused no apparent effects on RCR values, suggesting preserved OXPHOS coupling despite reduced complex I activity in the flight muscle (Figures 6 and 7A). In order to ensure that OXPHOS coupling was not affected, we then calculated two FCR values from PBS and Zymo-injected insects: the leak/ETS (L/E) and the OXPHOS/ETS (P/E) coupling control ratios. The L/E ratios reflect an indirect assessment of OXPHOS coupling through altered proton leak relative to the ETS respiratory rates and span from 0 (fully coupled) to 1 (fully uncoupled) values (Doerrier et al., 2018). Figure 7B shows that L/E ratios hardly changed upon Zymo injection over time indicating that mitochondrial inner membrane permeability to protons was not affected. The P/E ratios represent the limiting effect of the mitochondrial phosphorylation system, which comprises the ATP synthase, the phosphate carrier and the adenine nucleotide translocase, to respiratory rates relative to the ETS (Doerrier et al., 2018). P/E values span from minimal values of 0 (total inability to perform OXPHOS due to the maximal limitation of the phosphorylation system) to 1 (full OXPHOS capacity due to the maximal contribution of the phosphorylation system). Figure 7C shows that no significant changes were observed on P/E by Zymo injection. However, a significant effect of time (F_time_ *p* = 0.0004) at 12 h was identified regardless of the treatment, suggesting that activation of the innate immune response drives the mitochondrial phosphorylation system as an attempt to overcome reduced complex I activity (Figures 5 and 6). In this regard, a ∼150 fold increase in defensin A expression was observed in the PBS-injected insects (Figure 3D) suggesting that even pricking the insect with a sterile solution is enough to drive a compensatory mitochondrial metabolic response through mild activation of the innate immunity (Figure 7C).

**Figure 7.**
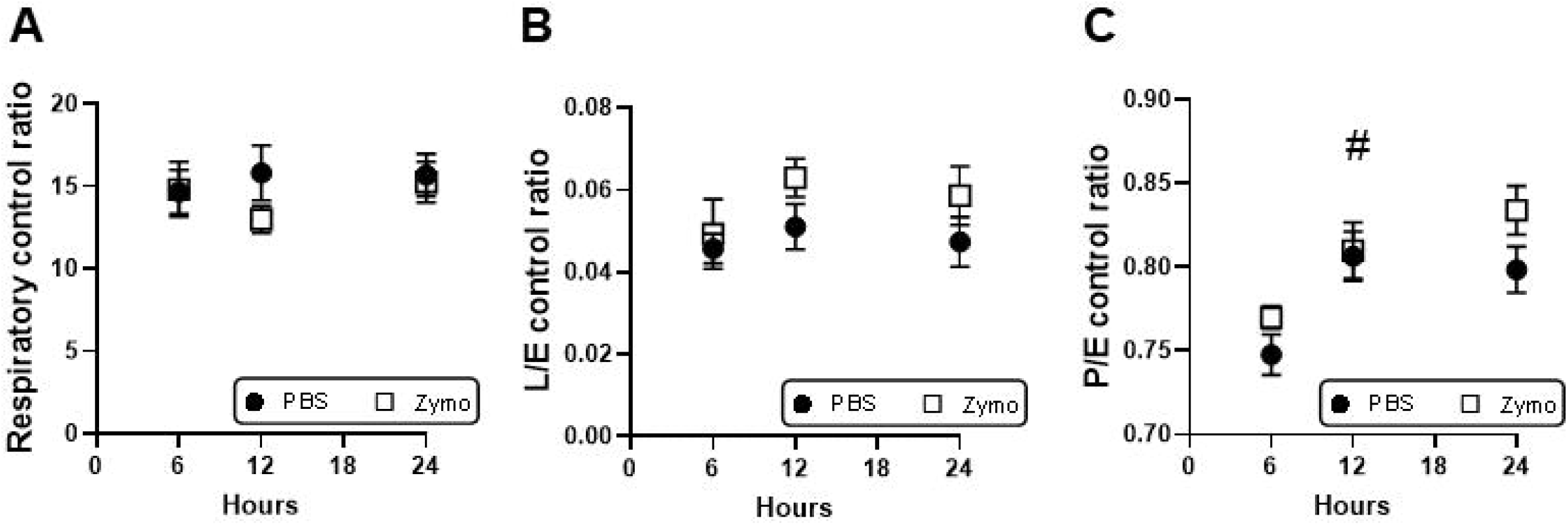
OXPHOS coupling efficiency is preserved despite reduced complex I activity. Respiratory control ratios (A) and flux control ratios (FCR, B and C) of permeabilized flight muscles from PBS (black circles) and Zymo (white squares) injected insects were quantified along 24 h by using 10 mM pyruvate-proline, 2 mM ADP and 2.5 µM FCCP following by the addition of 0.5 μM rotenone. (B) L/E ratios were calculated by dividing the leak O_2_ consumption rates (L) by their respective ETS respiratory rates (E). (C) P/E ratios were calculated by dividing the OXPHOS O_2_ consumption rates (P) by their respective ETS respiratory rates (E). Comparisons between groups were done by 2-way ANOVA and Šídák’s multiple comparisons test F_time_ (1.99, 23.9) = 11.25, *p* = 0.0004 # F_inject_ (1, 22) = 1.27, p = 0.273. Data are expressed as mean ± standard error of the mean (SEM) of at least four different experiments for both PBS and Zymo groups.

### 2.6. Reduced complex I activity by innate immune activation compromises mitochondrial proline oxidation

Mitochondrial proline oxidation represents a critical metabolic pathway to provide the energy required to sustain flight in many insect species, particularly blood-feeding insect vectors (Bursell, 1975; Scaraffia and Wells, 2003). To ensure that the reduced respiratory capacity observed in *A. aegypti* flight muscle upon Zymo injection was due to impaired proline oxidation, we assessed the O_2_ consumption rates of permeabilized flight muscles using only proline as a substrate. Experiments performed 24 h upon injections revealed that innate immune activation significantly reduced OXPHOS, ETS, and proline dehydrogenase-mediated respiratory rates (Figures 8B, 8D and 8E). Likewise, the experiments carried out using pyruvate and proline as substrates (Figure 6), we observed that RCR, L/E, and P/E values were not affected by Zymo suggesting that activation of innate immune response did not alter proline-fueled OXPHOS coupling efficiency (Figures 8F, 8G and 8H).

**Figure 8.**
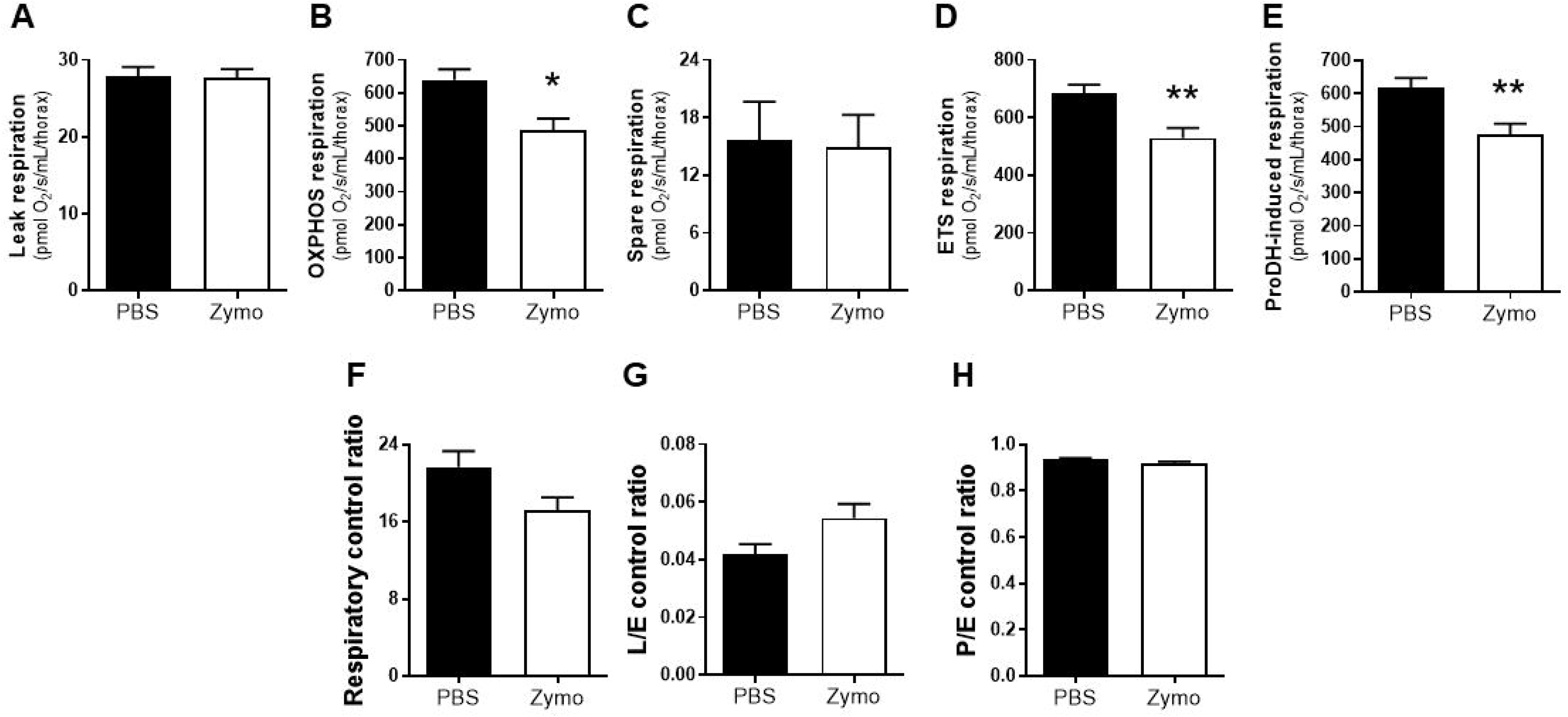
Innate immune activation targets mitochondrial proline oxidation. Respiratory rates of mechanically permeabilized *A. aegypti* flight muscle from PBS (black bars) or Zymo (white bars) were determined 24 h upon injections using 20 mM proline as a substrate. (A) Leak respiration. (B) OXPHOS respiratory rates. Comparisons between groups were done by Mann Whitney with * *p* = 0.014 relative to PBS. (C) Spare respiration. (D) ETS respiration. Comparisons between groups were done by Mann Whitney with * *p* = 0.0093 relative to PBS. (E) ProDH respiration. Comparisons between groups were done by Mann Whitney with * *p* = 0.014 relative to PBS. Data are expressed as mean O_2_ consumption rates (pmol/s/mL/thorax) ± standard error of the mean (SEM) of at least six different experiments.

### 2.7. The magnitude of innate immune response activation negatively correlates with flight muscle respiratory rates

Several lines of evidence indicate that in human sepsis, immune response activation has detrimental effects on energy metabolism, mitochondrial structure, and function (Brealey et al., 2004, 2002; Japiassú et al., 2011; Singer, 2014). Indeed, loss of complex I activity, increased proton leak and inhibition of the mitochondrial phosphorylation system are just some of the phenotypes so far reported on either experimental or clinical sepsis (Brealey et al., 2002; d’Avila et al., 2008; Japiassú et al., 2011). Importantly, the severity of sepsis strongly correlates with the degree of changes in mitochondrial processes and OXPHOS (Brealey et al., 2002; Japiassú et al., 2011; Sjövall et al., 2010). Given that our method to assess respiratory rates in *A. aegypti* can be carried out at the individual level, we correlated defensin A expression in the abdomen (fat body) of the very same individual where respiratory rates were assessed in the flight muscle (Gaviraghi et al., 2021; Gaviraghi and Oliveira, 2019). Figure 9 shows that the magnitude of innate immune activation, as measured by defensin A expression in the fat body, negatively correlate with flight muscle respiratory rates when using pyruvate and proline as substrates. This inhibitory effect was consistent and significant on OXPHOS, ETS, and rotenone-sensitive respiratory rates, strongly suggesting that factors released by the activation of innate immune response in the fat body have systemic effects that target flight muscle complex I function. Beyond OXPHOS, ETS and rotenone-sensitive respiration, no significant correlations were observed between defensin A expression and all other mitochondrial metabolic states in the flight muscle (Table 1). We conclude that the magnitude of complex I dysfunction as a result of the innate immune activation (Figure 9C) compromises not only the maximal respiratory capacity of flight muscle mitochondria (Figure 9B), but also their ability to synthesize ATP by the OXPHOS (Figure 9A).

**Table 1:** Correlation analyses between mitochondrial respiratory rates or flux control ratios and immune activation in *A.aegypti*.

| **Mitochondrial metabolic states** | **r** | **p** | **n** |
| --- | --- | --- | --- |
| 6h Leak respiration | -0.13 | 0.714 | 11 |
| 12h Leak respiration | 0,26 | 0,299 | 17 |
| 24h Leak respiration | - 0,15 | 0,478 | 24 |
| 6h L/E control ratio | -0.009 | 0.99 | 11 |
| 12h L/E control ratio | 0,34 | 0,178 | 17 |
| 24h L/E control ratio | 0,34 | 0,105 | 24 |
| 6h OXPHOS respiration | -0,14 | 0.698 | 11 |
| 12h OXPHOS respiration | - 0,34 | 0,171 | 17 |
| 6h P/E control ratio | 0.09 | 0.800 | 11 |
| 12h P/E control ratio | 0,28 | 0,501 | 17 |
| 24h P/E control ratio | 0,07 | 0,728 | 24 |
| 6h ETS respiration | -0.04 | 0.924 | 11 |
| 12h ETS respiration | - 0,34 | 0,188 | 17 |
| 6h Spare respiration | -0.15 | 0.673 | 11 |
| 12h Spare respiration | 0,04 | 0,868 | 17 |
| 24h Spare respiration | - 0,24 | 0,266 | 24 |
| 6h Rotenone-sensitive respiration (complex I) | -0.14 | 0.698 | 11 |
| 12h Rotenone-sensitive respiration (complex I) | - 0,34 | 0,188 | 17 |
| 6h Rotenone-resistant antimycin-sensitive respiration (Proline DH) | -0.09 | 0.796 | 11 |
| 12h Rotenone-resistant antimycin-sensitive respiration (Proline DH) | - 0,23 | 0,367 | 17 |
| 24h Rotenone-resistant antimycin-sensitive respiration (Proline DH) | - 0,33 | 0,110 | 24 |
| 6h Respiratory control ratio (RCR) | -0.04 | 0.924 | 11 |
| 12h Respiratory control ratio (RCR) | - 0,26 | 0,303 | 17 |
| 24h Respiratory control ratio (RCR) | - 0,28 | 0,181 | 24 |

Spearman r and p values were obtained from correlation analyses between the stated mitochondrial metabolic states and the levels of defensin a expression determined in adult females of *A. aegypti*. The total population of PBS and zymosan-injected insects (n=11, 17 and 24 at 6, 12 and 24h, respectively) were considered in the correlation analyses.

**Figure 9.**
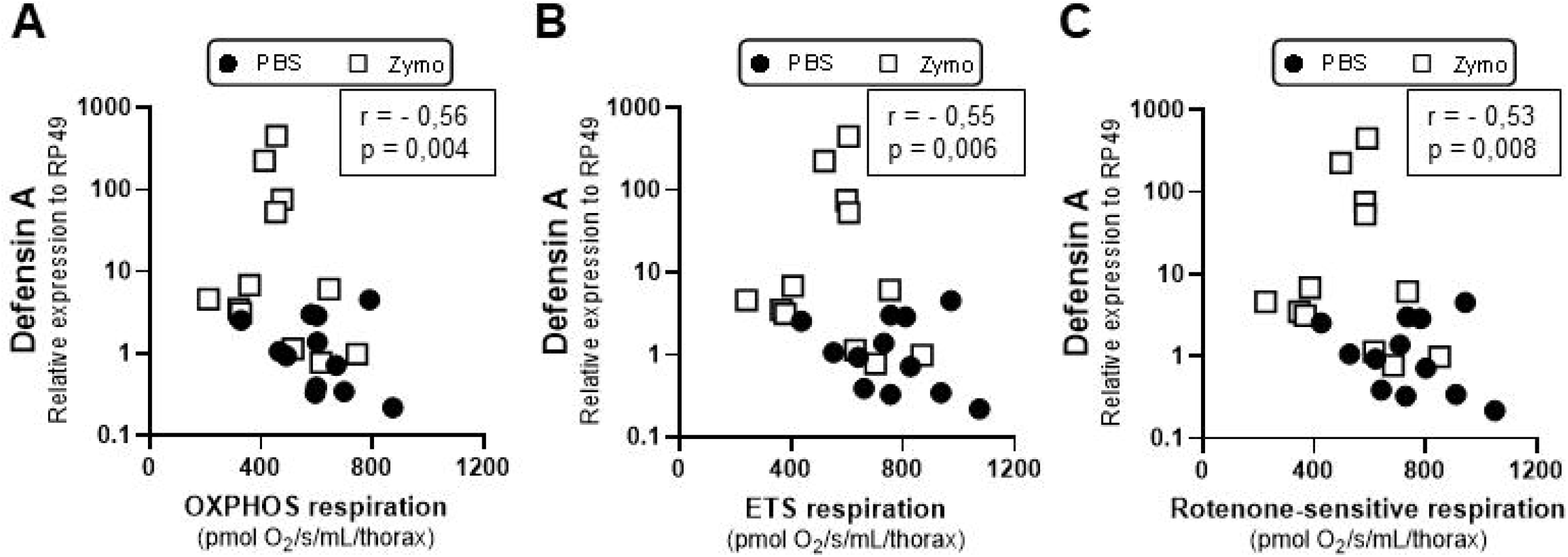
Respiratory rates negatively associate to the magnitude of innate immune activation. The activation of innate immune response measured as the defensin a expression levels in the fat body were correlated with their respective flight muscle respiratory rates linked to ATP synthesis (OXPHOS respiration, A), to the electron transport system (ETS respiration, B) or from complex I activity (Rotenone-sensitive respiration, C). Respiratory rates from PBS (black circles) and Zymo (white squares) exposed insects 24 h upon injection were compared using the Spearman r correlation tests. Every point represents an *A. aegypti* individual where defensin a expression in the fat body was matched with its respective respiratory rate in the flight muscle. Spearman r correlation and *p* values were - 0.56 and 0.004 (A), -0.55 and 0.006 (B), -0.53 and 0.008 (C).

## 3. Discussion

The energy costs posed by the activation of the host immune response are high and represent a critical challenge to avoid loss of energy homeostasis (Rivera et al., 1998; Tidball, 2005). Some of the metabolic outputs reported during innate immune response activation include shifts in nutrient utilization, insulin resistance, and mitochondrial dysfunction which are necessary to meet not only the cellular energy demands but also to allow cell survival (Rera et al., 2012; Shi et al., 2006; Singer, 2014). In several experimental and clinical models of inflammatory challenges, activation of innate immune response targets a variety of mitochondrial processes including the TCA cycle, mitochondrial membrane potential, ATP synthesis and oxidant production (d’Avila et al., 2008; Japiassú et al., 2011; Mainali et al., 2021; Stanzani et al., 2019). However, it remains largely obscure whether the metabolic alterations caused by the activation of innate immune response are evolutionarily conserved in invertebrate models (Chikka et al., 2016). This is particularly relevant considering the case of insect vectors of NTD, which are naturally exposed to numerous pathogens that cause important human diseases. Here demonstrate that sterile activation of innate immunity in the arbovirus vector *A. aegypti* by Zymo strongly reduced flight muscle complex I activity directly affecting OXPHOS respiratory capacity, mitochondrial proline oxidation, and flight activity. This effect does not involve modulation of mitochondrial content through mitophagy or reduced biogenesis, nor a broader effect on mitochondrial carriers, TCA cycle enzymes, or any other OXPHOS component. Rather, a selective effect on complex I activity was observed 24 h after Zymo injection. A general outline of the main results of the present work is given in Figure 10.

**Figure 10.**
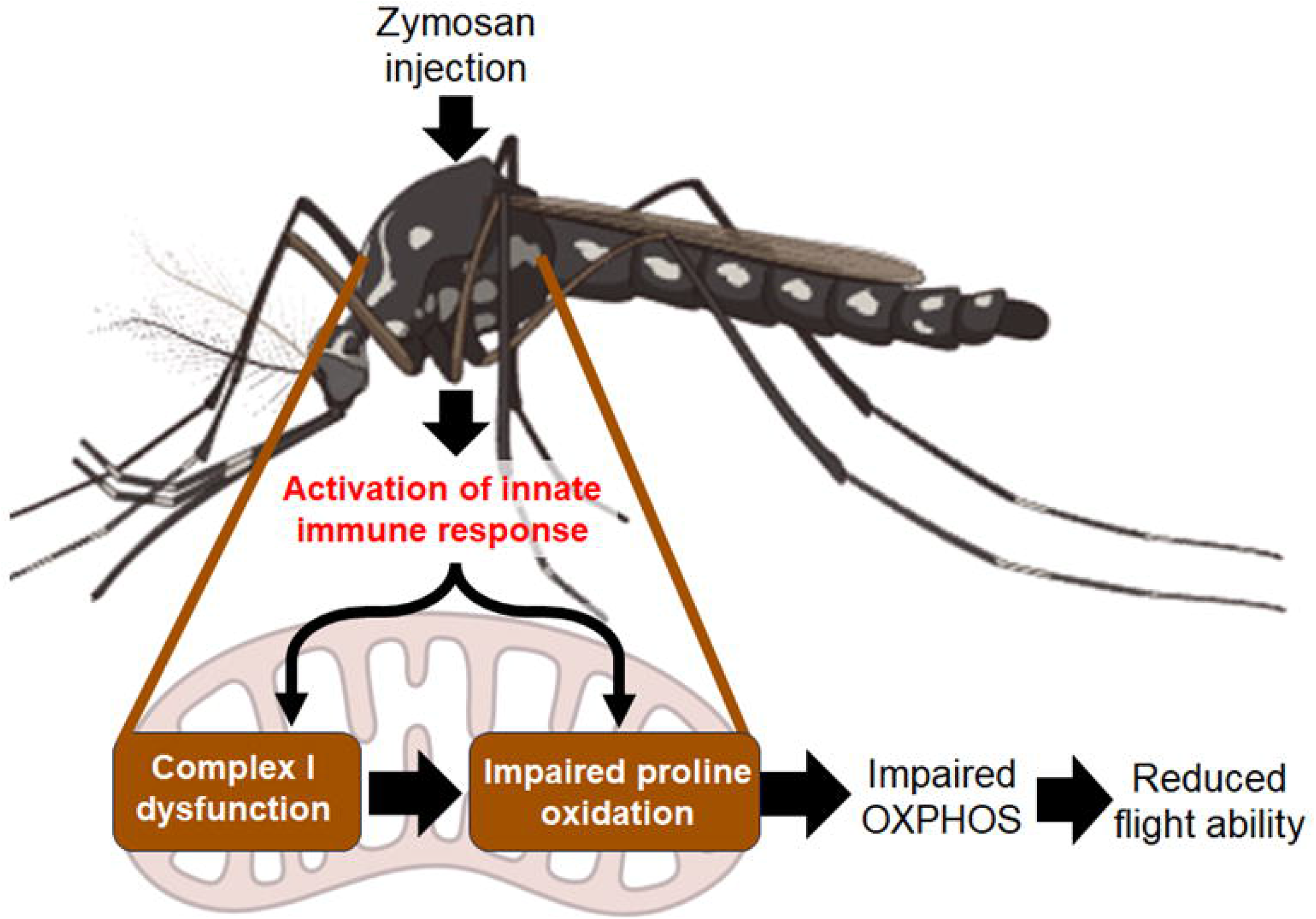
Schematic representation of the metabolic consequences to flight muscle mitochondria upon the activation of innate immune response in *A. aegypti*. The Upon zymosan injection a time-dependent activation of the innate immune response takes place specifically targeting mitochondrial flight muscle complex I activity. Despite this, OXPHOS coupling efficiency was not affected. However, mitochondrial oxidation of the main fuel to sustain flight activity (proline) is also compromised by the activation of innate immune response. Thus, the capacity of flight muscle to sustain maximal ATP synthesis is affected, which ultimately impairs induced flight ability in adult females. Further details are provided in the discussion section.

Despite the heterogeneity of metabolic responses in response to inflammation, reduced ETS complex activities seem a common phenotype observed in many models of innate immune activation (Brealey et al., 2004, 2002; Garaude et al., 2016; Merz et al., 2017; Vanasco et al., 2014). For example, in human sepsis complex I activity negatively correlates with nitric oxide (NO) levels, strongly associating with multiorgan dysfunction and poor patient outcome (Brealey et al., 2002). On the other hand, pharmacological and genetic loss of function experiments targeting complex I generate a potent pro-inflammatory phenotype which involves increased mitochondrial oxidant production (Cai et al., 2023; Groß et al., 2016; Jin et al., 2014). This suggests the existence of a reciprocal negative association between complex I activity and the intensity of immune response, but the underlying mechanisms remain to be elucidated. Indeed, we demonstrate that the intensity of innate immune activation (as measured by the expression of defensin a in the fat body) negatively correlates with flight muscle OXPHOS and complex I activity at the individual level (Figure 9). Thus, the extent of metabolic consequences upon immune activation as a result of specific inhibition of complex I would depend on the magnitude of systemic signals released by host cells upon a PAMP challenge. Supporting this hypothesis, our laboratory has previously reported that pharmacological inhibition of complex I in adult *A. aegypti* females directly impacted induced flight activity suggesting that preserved OXPHOS through complex I is essential for flight dispersal (Gaviraghi and Oliveira, 2020).

Although a direct mechanist explanation by which activation of innate immune response in *A. aegypti* drives complex I dysfunction remains to be elucidated, we think that some processes should be considered. The first one is the potential involvement of RNS such as NO and peroxynitrite (ONOO-) as mediators of complex I dysfunction. In fact, loss of complex I activity can be triggered by RNS during inflammatory challenges in mammalian models (Bailey et al., 2019; Brealey et al., 2002; Dahm et al., 2006; Murray et al., 2003; Riobó et al., 2001). NO is a classical signaling molecule released during inflammatory challenges, and targets several energy metabolism enzymes including AMP protein kinase (AMPK), as well as ETS complexes I and IV (Almeida et al., 2004; Cleeter et al., 1994; Dahm et al., 2006). On the other hand, ONOO- is produced from the spontaneous reaction of superoxide with NO, driving protein nitration and potent inactivation of several mitochondrial enzymes (Cassina and Radi, 1996; Castro et al., 1994; Murray et al., 2003; Radi et al., 1994). Considering that mitochondria represent the major site of cellular superoxide production (Boveris et al., 1972; Boveris and Chance, 1973), it seems plausible that ONOO-production would be favored in this organelle. In fact, mitochondria represent not only a source of endogenous ONOO-production (Valez et al., 2013) but is also the major site of nitrated proteins (Sacksteder et al., 2006). Indeed, some of the previously reported mitochondrial effects attributed to NO seem to be caused by ONOO-(Dahm et al., 2006). While NO interacts mostly to cytochrome *c* oxidase through a loose and reversible mechanism (Cassina and Radi, 1996; Cleeter et al., 1994), mitochondrial protein oxidation and nitration by ONOO-irreversibly impair key mitochondrial enzyme activities (Cassina and Radi, 1996; Castro et al., 1994; Radi, 2013; Radi et al., 1994). Despite NO and ONOO- mediate bioenergetic remodeling, we think that specific reduction of complex I activity by immune activation in *A. aegypti* does not involve RNS metabolism for the following reasons: *i) A. aegypti* NOS expression was not altered upon Zymo injection, suggesting that NO production would not be activated in our settings (Figure 3C); *ii)* complex I driven-respiration was only affected at 24 h upon Zymo injection, suggesting that mitochondrial effects by immune response activation take place at later stages (Figures 5D and 5F). One might think that NOS activation by Zymo would be even earlier than 6 h which would drive delayed and persistent complex I dysfunction. If that was the case, then respiratory rates mediated by complex I would be lowered before 24 h, a condition not supported by our results (Figure 5); *iii)* total oxidant levels were not altered 24 h after Zymo injection, strongly indicating that high superoxide and NO levels required to allow ONOO- production were not present in our experimental setting by the time of complex I dysfunction (Figure 4). Finally, as NO drives mitochondrial superoxide production in mammalian models (Riobó et al., 2001), the absence of redox imbalance upon Zymo injection strengthens our view that reduced complex I activity does not involve RNS metabolism in adult *A. aegypti*.

The second aspect considers the possibility that defensin A, and potentially other antimicrobial peptides (AMPs) would directly interact with and modulate mitochondrial processes. In this regard, PAMPs release upon pathogen infection in host cells mediate the expression of AMP by activating specific signaling cascades. AMPs are cationic and amphipatic molecules that mediate pathogen killing in vertebrates and insects (Wu et al., 2018). Considering the similarities between bacterial and mitochondrial membranes, the amphipathic nature of AMPs, and their selectivity towards bacterial membranes, it is possible that AMPs would directly interact with mitochondria (Karlin et al., 1999; Yeaman and Yount, 2003). Supporting evidence demonstrates that in several organism models AMPs promote redox imbalance, depolarization of mitochondrial membrane potential, cytochrome *c* release, and the induction of a pro-apoptotic program resulting in cell death (Cerón et al., 2010; Cho et al., 2012; Hwang et al., 2011). Interestingly, the *Drosophila* AMP metchnikowin targets fungi complex II activity strengthening that mitochondrial physiology and energy metabolism are affected by AMPs (Moghaddam et al., 2017). The overexpression of *relish,* a transcription factor required for AMP expression in *Drosophila*, promoted mitochondrial membrane potential depolarization and apoptosis in fat body cells and reduced longevity (Badinloo et al., 2018). In *A. aegypti*, Zymo activates the innate immune response through Toll, IMD and Jak/STAT signaling pathways upregulating AMP expression (Barletta et al., 2012). Although the depolarization of mitochondrial membrane potential is a metabolic hallmark of AMP, a condition not observed in our experimental conditions (Figure 5C and 8A, Table 1), we think plausible that reduced complex I activity would be driven by the following possibilities: *i)* a direct interaction of defensin A on complex I; *ii)* an indirect effect of an immune response signaling cascade targeting complex I. In this regard, evidence from mammals indicates that Ecsit protein, a cytosolic mediator of Toll pathway, is required for complex I assembly and stability suggesting a direct link between immune activation and mitochondrial metabolism (Vogel et al., 2007). Conceivably, *A. aegypti* complex I structure is somehow distinct from other organisms rendering it more susceptible to immune signals. In this sense, *in silico* evidence on KEGG database indicates that *A. aegypti* codes only 42 out of 45 human complex I subunits (https://www.genome.jp/kegg-bin/show_pathway?aag00190). The missing subunits belong to the supernumerary transmembrane helices (*Ndufa3*, and *Ndufc1*) and to the hydrophilic matrix arm (*Ndufv3*) (Zhu et al., 2016). Interestingly, the absence of the same subunits is shared by *D. melanogaster* suggesting that a smaller complex I is a feature of insect mitochondria (https://www.genome.jp/pathway/dme00190+M00154). In any case, whether the absence of the three subunits would facilitate direct interaction of defensin A to insect complex I, or rendering it more susceptible to an immune response signaling cascade, remains to be elucidated.

Despite the specific effect of innate immune activation by Zymo injection on *A. aegypti* complex I activity, we observed that OXPHOS coupling and efficiency were preserved (Figures 7 and 8). This strongly suggests that proton leak through the mitochondrial inner membrane was not stimulated due to its loss of stability and integrity by immune response activation. Rather, the mechanisms involved on mitochondrial electron supply to maintain OXPHOS were targeted by immune activation. Thus, reduced electron delivery through complex I seems to affect mitochondrial ATP output, OXPHOS, energy homeostasis and ultimately flight ability. In this sense, evidence demonstrates that a novel member of the SLC25A mitochondrial carrier family in *Anopheles gambiae* (AgMC1) revealed its contribution to the maintenance of mitochondrial membrane potential, oxidant production and determining the susceptibility to *Plasmodium* infections (Gonçalves et al., 2012). This strongly indicates that mechanisms aimed to provide mitochondrial electron supply regulate the susceptibility to a pathogen infection in an insect vector. In this regard, the establishment of a resistant phenotype against *Plasmodium* infection involve reduced OXPHOS and increased mitochondrial oxidant production indicating that regulation of mitochondrial energy metabolism control the susceptibility of insect vector to infection (Oliveira et al., 2011). In addition, dynamic modulation of metabolic enzymes during infections was reported during mosquito infection suggesting that bioenergetic mechanisms that supply electrons to OXPHOS play a role in the adaptive vector response against infectious challenges (Martins et al., 2021; Santana-Román et al., 2021; Vasconcellos et al., 2022, 2020).

The results observed in the present work raise the question on what are the metabolic consequences to flight dispersal of an infected *A. aegypti* female. In this sense, the dispersal capacity represents a key parameter not only for egg laying but also for pathogen transmission by insect vectors (Kaufmann et al., 2013; Petrić et al., 2014). To accomplish such an extremely energy-costing biological task, flight activity is essentially sustained by OXPHOS which take place within the huge mitochondria found in insect flight muscles (Mesquita et al., 2021). Our laboratory has investigated several mitochondrial processes in *A. aegypti* and revealed that flight muscle respiratory rates and mitochondrial oxidant production are sustained by proline, pyruvate and G3P oxidation (Gaviraghi et al., 2019; Gaviraghi and Oliveira, 2019; Soares et al., 2015). Mitochondrial oxidation of different substrates promoted distinct bioenergetic capacities but preserved OXPHOS efficiencies, underscoring the importance of mitochondrial electron supply for flight muscle energy homeostasis. We also identified that pharmacological inhibition of complex I directly affected induced flight activity, strengthening that suitable mitochondrial electron supply is of central importance to insect dispersal (Gaviraghi and Oliveira, 2020). However, direct observations associating innate immune activation and loss of energy homeostasis in insect vectors remain largely unknown. Mitochondrial proline oxidation is a key substrate for OXPHOS and fuels *A. aegypti* flight activity (Gaviraghi and Oliveira, 2019; Scaraffia and Wells, 2003; Soares et al., 2015). Since proline oxidation generates L-ketoglutarate for the TCA cycle, a functional complex I is required to provide matrix NAD+ for the dehydrogenases involved in this metabolic pathway. Indeed, our results indicate that mitochondrial proline oxidation was suppressed upon Zymo injection, directly impacting respiratory rates but not OXPHOS coupling (Figure 8). Importantly, impaired proline oxidation might explain the reduced flight activity observed in Zymo-injected insects (Figure 3E).

From the evolutive perspective, functional associations between complex I and immunity were reported for invertebrate organisms (Chikka et al., 2016; Cormier et al., 2022; Mello et al., 2022). In this sense, specific complex I-driven OXPHOS associates with increasing defensin expression in honeybees during winter (Cormier et al., 2022). Remarkably, a shift towards G3P and succinate oxidation was observed suggesting that functional suppression complex I-dependent OXPHOS is linked to immune activation (Cormier et al., 2022). In *C. elegans*, while strong inhibition of complex I activates innate immune response and prevents neurodegeneration (Chikka et al., 2016), mild rotenone treatment down-regulated the expression of immune response genes, rendering roundworms prone to bacterial infections (Mello et al., 2022).

In summary, we demonstrate that activation of innate immunity is strongly associated with reduced flight muscle complex I activity with direct consequences to mitochondrial proline oxidation and flight activity. Conceivably, the fitness costs associated with reduced complex I activity in the flight muscle would limit the huge energy required to sustain muscle contraction and the high wing beat frequencies in mosquitoes (Bomphrey et al., 2017). Remarkably, our results indicate that a trade-off between dispersal and immunity exists in an insect vector, underscoring the potential consequences of disrupted flight muscle mitochondrial energy metabolism to arbovirus transmission.

## 4. Experimental procedures

### 4.1. Reagents

ADP was purchased from Selleckchem (cat. N° S6325). Sucrose (cat. N° S8501), antimycin a (AA) (cat. N° A8674), zymosan (cat. N° Z4250), KCl (cat. N° P9541), KH_2_PO_4_ (cat. N° P5655), HEPES (cat. N° H7006), EGTA (cat. N° E3889), Sodium ascorbate (cat. N° A7631), dihydroethidium (cat. N° D7008), BSA fatty acid-free (cat. N° A6003), Pyruvate (cat. N° P2256), Proline (cat. N° P0380), carbonyl cyanide p-(trifluoromethoxy) phenylhydrazone (FCCP) (cat. N° C2920), rotenone (cat. N° R8875), Sucrose (cat. N° S8501), Tris (cat. N° T1699), Acetyl-CoA (cat. N° A2056), DTNB (cat. N° D8130), Oxaloacetate (cat. N° O4126), Cytochrome c (cat. N° C2506) were from Sigma-Aldrich. TRIzol reagent was purchased from ThermoFisher Scientific, and DNAse I from Fermentas. High-capacity cDNA reverse transcription kit, SYBR Green PCR Master mix, StepOnePlus Real Time PCR System were from Applied Biosystems. MgCl_2_ (cat. N° V000149) was from Vetec, KCN (cat. N° 1049670100) from Supelco, and TMPD (cat. N° 87890) was from Fluka. NADH (cat. N° 16078) and Glycerol-3-Phosphate (cat. N° 20729) were from Cayman. Schneideŕs *Drosophila* medium (cat. N° 21720024) was from Gibco. ADP, pyruvate, ascorbate, proline, cytochrome c, KCN, NADH, glycerol 3 phosphate, acetyl-CoA, oxaloacetate, TMPD, were dissolved in deionized H_2_O. For ADP the pH was adjusted to pH 7.2 with NaOH. AA, FCCP, and rotenone were dissolved in ethanol p.a. All solutions were aliquoted and stored at -20°C, except pyruvate, ascorbate, TMPD, KCN which were dissolved in deionized H_2_O fresh on the day of each experiment.

### 4.2. Ethics statement

All experimental protocols and animal care were carried out in accordance to the institutional care and use committee (Comitê para Experimentação e Uso de Animais da Universidade Federal do Rio de Janeiro/CEUA-UFRJ) and the NIH Guide for the Care and Use of Laboratory Animals (ISBN 0–309-05377-3). The protocols were approved under the registry CEUA-UFRJ #155/13.

### 4.3. Insects

*Aedes* (Stegomyia) *aegypti* (Linnaeus, 1762), red eyes strain, were maintained at 28°C, 70– 80% relative humidity with a photoperiod of 12 h light/dark (L:D, 12:12 h). Larvae were reared on a diet consisting of commercial dog chow. Usually about 200 insects were placed in 5 L plastic cages in a 1:1 sex-ratio and allowed to feed *ad libitum* on cotton pads soaked with 10 % (w/v) sucrose solution. For all experiments, only female mosquitoes with 5-7 days after the emergence were used.

### 4.4. Activation of immune response

To induce systemic immune activation, we challenged female mosquitoes by pricking their thorax with a glass needle (3.5” Drummond, Drummond Scientific, Broomall, PA, USA) embedded with either PBS or 5 mg/mL zymosan A solution. Mosquitoes were dissected for RNA extraction and respirometry at 6, 12 and 24 h after the immune challenge. Non-injected insects revealed to be essentially identical to PBS-injected ones in all parameters assessed and for this reason were not included in the figures (data not shown). Zymosan A solution was prepared according to manufacturer’s instructions. Briefly, a suspension of zymosan/PBS was autoclaved before use to avoid the growth of any potential live yeast contaminants.

### 4.5. RNA extraction and quantitative PCR

Total RNA from mosquito fat body was extracted using the TRIZOL reagent following the manufacturer’s instructions. RNA was treated with DNAse I and first-strand cDNA synthesis were carried out using High-Capacity cDNA Reverse transcription kit. The amplification efficiency of the experimental set for each gene was tested with serial dilutions of cDNA and was only used if the resultant efficiency was ≥ 90%. Each PCR reaction (15 μL volume) contained diluted cDNA, Power SYBR Green PCR Master Mix and 300 nM of forward and reverse primers. Quantitative PCR was performed in a StepOnePlus Real Time PCR System using Applied Biosystems recommended qPCR conditions (20 seconds at 95°C followed by 40 cycles of 95°C for 1 second and 20 seconds at 60°C followed by a melting curve to assure a single product was amplified). The comparative ΔΔCt method was used to evaluate changes in gene expression levels and all standard errors were calculated based on ΔCt as described in Applied Biosystems User Bulletin #2 (http://www3.appliedbiosystems.com/cms/groups/mcb_support/documents/generaldocuments/cms_040980.pdf). *A. aegypti* ribosomal protein 49 gene, RP-49, was used as endogenous control (accession number AAT45939), as described elsewhere (Gentile et al., 2005). Each figure represents at least five biological replicates with three technical replicates for each sample. Specific qPCR primers used in this study were the following: Defensin a (accession number AAEL003841) forward: GATTCGGCGTTGGTGATAGT and reverse: TTATTCAATTCCGGCAGACG; Nitric oxide synthase (accession number AAEL009745) forward: GCCAATTTGGACGGAGAGGA and reverse: CGATCGGAACGGTGCAATTC; RP49 (accession number AAEL003396) forward: GCTATGACAAGCTTGCCCCCA and reverse: TCATCAGCACCTCCAGCT.

### 4.6. Lifespan

To determine the lifespan, after injection with Zymosan A or PBS, dead mosquitoes were removed by suction and recorded every day for seven days.

### 4.7. Respirometry analyses

The respiratory activity of mechanically permeabilized flight muscles from *A. aegypti* was carried out in a two-channel respirometer (Oxygraph-2k, O2k Oroboros Instruments, Innsbruck, Austria) at 27.5°C and under continuous stirring set up at 750 rpm as previously established by our group (Gaviraghi et al., 2021; Gaviraghi and Oliveira, 2019; Soares et al., 2015). A single cracked thorax was added to the O2k chamber filled with "respiration buffer" (120 mM KCl, 5 mM KH_2_PO_4_, 3 mM Hepes, 1 mM EGTA, 1.5 mM MgCl_2_, and 0.2 % fatty acid free bovine serum albumin, pH 7.2) and allowed to equilibrate for about 5 min. During this time, flight muscle was mechanically permeabilized by the magnetic stirring of the respirometer chamber. At the same time, an appropriate amount (5–10 mL) of an O_2_-enriched gaseous mixture (70% O_2_ and 30% N_2_ mol/mol) was injected through the stopper of each chamber using a 60 mL syringe to reach an O_2_ concentration close to 500 nmol/mL. This step was introduced to avoid potential effects of O_2_ diffusion and electron transfer due to O_2_ shortage during measurements (Pesta and Gnaiger, 2012). Also, injections of O_2_-enriched gaseous mixture were performed once the O_2_ concentration fell down below 150 nmol/mL into the O2k-chamber. Mitochondrial physiology was assessed by high resolution respirometry coupled to substrate-uncoupler-inhibitor-titration (HRR-SUIT) assay to establish RCR and FCR as described in the literature (Doerrier et al., 2018). A general outline of a typical HRR experiment is depicted in figure 5A. Two distinct oxidizable substrate combinations were used in our experiments: pyruvate + proline (Pyr+Pro) or only proline (Pro). The experiment was started by the addition of saturating concentrations of substrates (10 mM pyruvate + 10 mM proline or 20 mM proline alone). The leak respiratory rates (Leak) are characterized by a non-phosphorylating condition with high concentration of oxidizable substrates, but in the absence of ADP, and was calculated by subtracting the O_2_ consumption rates after addition of 2.5 µg/mL AA from the respiratory rates induced by oxidizable substrates alone. Subsequently, the synthesis of ATP was induced by 1 mM ADP followed by a second shot reaching the final concentration of 2 mM. The O_2_ consumption coupled with oxidative phosphorylation (OXPHOS) respiratory rates were calculated by subtracting the ADP-stimulated O_2_ consumption rates from the respiratory rates induced by the leak state. To assess maximal uncoupled respiratory rates (ETS) pharmacological uncoupling was induced by stepwise titration with carbonyl cyanide p-(trifluoromethoxy) phenylhydrazone (FCCP) to reach final concentrations of 2.5 µM and then subtracting respiratory rates by the residual O_2_ consumption after addition of 2.5 µg/mL AA. The spare respiratory rates were calculated by subtracting the ET capacity rates from the OXPHOS capacity. The contribution of complex I to respiratory rates was assessed by calculating the inhibitory effect of 0.5 μM rotenone on FCCP-uncoupled O_2_ consumption using 10 mM Pyr + Pro as substrates. Complex I-linked respiratory rates were calculated as the FCCP-uncoupled Pyr+Pro-induced O_2_ consumption rates subtracted by the rotenone-insensitive rates. The contribution of proline dehydrogenase (ProDH) to respiratory rates was assessed by calculating the inhibitory effect of stepwise titration of the ProDH inhibitor L-thiazolidine-4-carboxylic acid (T4C) (Elthon and Stewart, 1984; Magdaleno et al., 2009) up to 1 mM on FCCP-uncoupled O_2_ consumption using 20 mM Pro as substrate. ProDH-linked respiratory rates were calculated as the FCCP-uncoupled 20 mM Pro-induced O_2_ consumption rates subtracted by the T4C-insensitive rates. Finally, for all substrates, respiratory rates were inhibited by the addition of AA and the residual O_2_ consumption was calculated as the AA-insensitive rates.

### 4.8. Mitochondrial isolation

Isolation of mitochondria from flight muscle was performed as previously described by our group (Soares et al., 2015) with minor modification. About 24 mosquitoes were immobilized by chilling on ice, and then gently homogenized in a 3 mL glass tissue grinder with 2 mL of ice-cold isolation medium (250 mM sucrose, 5 mM Tris-HCl, 2 mM EGTA, pH 7.4) without fatty acid free bovine serum albumin. The preparation was maintained at 4°C throughout the subsequent washing and centrifugation procedures. The liquid was centrifuged at 300 x g for 5 min in an Eppendorf centrifuge model 5804-R with a rotor F45-30-11. The supernatant was collected and further centrifuged at 10,000 x g for 10 min. The brown pellet was carefully re-suspended in approximately 50 µL of "hypotonic buffer" (25 mM potassium phosphate buffer and 5 mM MgCl_2_, pH 7.2). Protein concentration was determined by the Lowry method, using bovine serum albumin as standard (Lowry et al., 1951).

### 4.9. Mitochondrial enzyme activities

Enzyme activities were determined in preparations of isolated mitochondrial from *A. aegypti* females following methods described in the literature (Kirby et al., 2007) with modifications. The mitochondrial preparation was subjected to three freeze-thawing cycles and the enzyme activities were determined spectrophotometrically using a Shimadzu spectrophotometer model visible 2450 (Shimadzu Scientific Instruments, Tokyo, Japan) at room temperature. Citrate synthase (CS) activity was determined by incubating a sample corresponding to 17 μg of protein from freeze-thawed mitochondria, in 75 mM Tris-HCl pH 8.0, 0.03 mM acetyl-CoA and 0.25 mM 5,5′-dithiobis (2-nitrobenzoic acid) (DTNB). The rate of reduced coenzyme A (CoASH) production was determined using the thiol reagent DTNB, which has an absorption maximum at 412 nm. The reaction was started by the addition of 0.5 mM oxaloacetate. CS activity was calculated by subtracting the rates of DTNB reduction induced by oxaloacetate from the basal using the molar extinction coefficient (ε = 13,600 M^−1^·cm^−1^) and expressed as nmol of reduced DTNB/min/thorax. Complex I activity of the ETS was determined as NADH:Cytc oxido-reductase activity by measuring the increase in absorbance at 550 nm due to the ferricytochrome c reduction. The reaction containing 1 mL of 100 mM potassium phosphate buffer, pH 7.4, 50 μM oxidized cytochrome *c*, 10 μg of protein from freeze-thawed mitochondria and 1 mM KCN was initiated by adding 200 μM NADH and the increase in absorbance at 550 nm was monitored for about 5 minutes. Rotenone (0.5 μM) was added to inhibit complex I activity, which was considered as the rotenone-sensitive rate of cytochrome c reduction (ε = 18.7 mM^−1^·cm^−1^). Glycerol 3-phosphate dehydrogenase (G3PDH) activity was determined as G3PDH:Cytc oxido-reductase and was measured in the same conditions described for NADH:Cytc oxido-reductase activity, but using 20 mM of sn-glycerol 3 phosphate (G3P) as substrate. Before substrate addition, 0.5 µM rotenone was added to avoid electron backflow from G3PDH to complex I. 2.5 μg/mL AA was added to inhibit complex III activity and the activity of G3P:Cytc oxido-reductase was considered as the AA-sensitive rate of cytochrome c reduction (ε = 18.7 mM^−1^·cm^−1^). Cytochrome c oxidase (COX) activity was measured polarographically at the end of the routine of respiratory analysis after AA using 2 mM ascorbate and 0.5 mM N,N,N’,N’-Tetramethyl-p-phenylenediaminedihydrochloride (TMPD), as an electron-donor regenerating system. To distinguish the O_2_ consumption deriving from the activity of COX from that due to TMPD chemical auto-oxidation, 5 mM KCN was added at the end of each experiment, and COX activity was considered as the cyanide-sensitive rate of O_2_ consumption. Since COX activity is limited by low O_2_ concentrations (Chandel et al., 1996), the O_2_ enriched gas mixture was also injected before measuring enzyme activity.

### 4.10. Induced flight activity test

To evaluate the effect of innate immune activation on mosquito flight capacity, we used the INduced FLight Activity TEst (INFLATE) developed and validated by our laboratory (Gaviraghi and Oliveira, 2020).

### 4.11. Intracellular oxidant quantification

The levels of intracellular oxidants were assessed in flight muscle from hemi-thorax of *A. aegypti*. Mosquitoes were anesthetized on ice bath and thoraces were dissected in Schneider’s Insect medium (SIM). Hemi-thoraces were incubated with 100 µM dihydroethidium (DHE) in SIM in darkness for 20 minutes at room temperature. After that, muscles were washed twice in SIM and hemi-thoraces fluorescence images were captured in a stereo microscope (Olympus model MVX10) using excitation at 546 nm and emission at 590 nm. Images analyzes were carried out in Image J/Fiji software (Schindelin et al., 2012) using only the red channel. Quantification of DHE in muscles was performed applying masks to define the area of interest and eliminating background signal. Average fluorescence intensity of masked images was registered and expressed as arbitrary fluorescence units.

### 4.12. Data and statistics

Data in graphs were presented as meanL±Lstandard error of the mean (SEM) of values for each condition. D’Agostino and Pearson normality tests were done for all values to assess their Gaussian distribution. If normality was achieved, then Student’s t-tests were performed for pairwise comparisons. If data did not follow normal distribution, then Mann-Whitney’s tests were performed for pairwise comparisons. For multiple comparisons between groups, Friedmańs and Durbin-Conover or 2-way ANOVA and *a posteriori* Šídák’s tests were carried out. Correlations between O_2_ consumption rates and defensin a expression levels were done by Spearman’s correlation test. Differences with p<0.05 were considered significant. Graphs were prepared by using the GraphPad Prism software version 6.00 for Windows (GraphPad Software, USA).

## Acknowledgements

We thank Mrs. Jaciara Miranda Freire for the excellent technical assistance on maintenance of *A. aegypti* colony. We also thank BSc. Carolina Chaves do Nascimento for the valuable help on statistical analyses. This study was financed in part by the Coordenação de Aperfeiçoamento de Pessoal de Nível Superior – Brasil (CAPES) – Finance Code 001, by the Conselho Nacional de Desenvolvimento Cientifico e Tecnológico (CNPq) [grant number: 404153/2016-0 MFO, and 483334/2013-8 AG], and the Fundação Carlos Chagas Filho de Amparo à Pesquisa do Estado do Rio de Janeiro (FAPERJ) [grant numbers E-26/102.333/2013, E-26/203.043/2016, and E-26/111.169/2011]. AG and MFO are fellows from CNPq [fellowship numbers 402409/2012-4 AG, and 303044/2017-9 and 308629/2021-3 MFO] and from PAPD-FAPERJ [fellowship number E-44/208702/2014]. The funders had no role in study design, data collection and analysis, decision to publish, or preparation of the manuscript.

## Author contributions

AG and MFO designed the work; ABFB and TLAS performed zymosan injections and qRT-PCR analyses. AG performed longevity, flight activity, mitochondrial O_2_ consumption and enzyme activities. MPO performed oxidant production by fluorescence microscopy. MHFS conceived and designed the innate immune experiments. AG and MFO wrote the manuscript. All authors critically revised the article and approved the final version of the manuscript.

## Abbreviated summary

- Innate immune activation impairs flight muscle complex I and flight ability.
- Reduced complex I compromise mitochondrial proline oxidation.
- The magnitude of innate immunity activation negatively correlates with complex I dysfunction.

## Abbreviations

G3PDH: Glycerol 3-phosphate dehydrogenase
DTNB: 5,5′-dithiobis 2-nitrobenzoic acid
T4C: L-thiazolidine-4-carboxylic acid
ATP: adenosine triphosphate
ADP: adenosine diphosphate
RCR: Respiratory control ratio
FCR: Flux control ratio
G3PDH: glycerol 3 phosphate dehydrogenase
CS: citrate synthase
COX: cytochrome *c* oxidase
HRR: high resolution respirometry
Zymo: zymosan
(SUIT): substrate-uncoupler-inhibitor titration
(DHE): dihydroethidium
(INFLATE): induced flight activity test
(ETS): electron transfer system
(OXPHOS): oxidative phosphorylation
(TOR): Target Of Rapamycin
(AMP): antimicrobial peptide
PAMP: Pathogen-associated molecular pattern
Pyr: pyruvate
Pro: proline
G3P: sn-glycerol 3-phosphate
FCCP: carbonylcyanide p-(trifluoromethoxy) phenylhydrazone
Rot: rotenone
AA: antimycin A
Asc: ascorbate
TMPD: N,N,N’,N,- Tetramethyl-p-phenylenediamine
KCN: potassium cyanide
oligo: oligomycin.

## Data availability statement

The data that support the findings of this study are available from the corresponding author upon request.

